# Versatile Method to measure locomotion

**DOI:** 10.1101/2020.03.20.000828

**Authors:** Taylor Barwell, Sehaj Raina, Laurent Seroude

**Affiliations:** Department of Biology, Queen’s University, BioSciences Complex, Kingston, Ontario K7L 3N6, Canada

**Keywords:** Behavior, FlyTracker, MatLab, aging, Drosophila melanogaster

## Abstract

Many studies require the ability to quantify locomotor behavior over time. The list of tracking softwares and their capabilities are constantly growing. At the 2019 CanFly Conference we presented preliminary results from an investigation of the effects of expressing polyglutamine repeats in fly muscles on longevity, locomotion and protein aggregation and received a lot of inquiries about our protocol to measure locomotion and how to use the FlyTracker MatLab software. This report describes a versatile locomotion measuring device and custom MatLab scripts for the extraction and analysis, and compilation of FlyTracker data in a format compatible with spreadsheet softwares. The measurement and analysis of multiple genotypes and both sexes across age shows that this method yields reproducible results that confirm that normal aging is associated with a progressive decline in locomotion as indicated by increased immobility and reduced velocity.

## Introduction

Locomotion is a fundamental behavioral trait involved directly or indirectly with almost all simple or complex behavior activities. It is also integral to studying a wide span of different biological phenomenon such as neurological or muscular pathologies, age associated changes and physiological responses. Measuring locomotion is challenging because of the many parameters that need to be controlled [1]. Additionally, the obtainment of large data sets requires automated system for tracking, data extraction and analysis.

Drosophila have proven to be a useful model to study some of the most complex behavioral phenotypes including: learning, memory [2, 3], sleep, circadian rhythms [4], courting, mating [5] and response to drugs, addiction [6]. With advances in technology numerous experimental setups, manual and automated tracking have been developed to describe and quantify fly locomotion [1].

Automated tracking relies on softwares that use a digital subtraction method to remove background and be able to distinguish the fly from optical or background artifacts [7-12]. Softwares can be used to track several flies at once in two or three dimensions, quantify specific fly behaviors (grooming, fighting, mating) as well as monitoring leg coordination of single animals [7, 12-19].

We chose the widely used, multi-platform FlyTracker MatLab software [15, 17, 20] that does not require any expensive equipment. Essentially anyone with any camera and a computer can run the software and perform these experiments. FlyTracker detects very accurately multiple flies at once in a video and is able to track the position, orientation and angle of the wings and legs, as well as distance between the fly and the wall of the chamber housing the fly. At the 2019 CanFly Conference we presented preliminary results from an investigation of the effects of expressing polyglutamine repeats in fly muscles on longevity, locomotion and protein aggregation. We received a lot of inquiries about our protocol to measure locomotion and how to use FlyTracker. Here, we report a locomotion measuring device that can accommodate 5 to 50 flies and custom MatLab scripts for the extraction and analysis, and compilation of FlyTracker data in a format compatible with spreadsheet softwares. Multiple genotypes, both sex and two experimental replicates across age demonstrate the reproducibility achieved.

## Materials and Methods

### Construction of the arena

A drill press was used to drill 2.5cm circles into thin (3mm), clear plexi glass in rows of 5. The rows were cut into strips (Figure 1A). A piece of clear plexi glass the same dimensions as the strip was adhered to the back of the strip with superglue. By constructing chambers in strips the arena can be adapted to accommodate experiments requiring 5 chambers and up to 50. A base was made from another piece of plexi glass, large enough to fit all 10 strips (Figure 1A). A small hole is drilled near the edge of the base to load the flies. Three small pieces of plexi glass were cut and glued to the base to use as a guide for positioning the strips consistently. Legs were made for the base out of cuvettes.

**Fig. 1.**
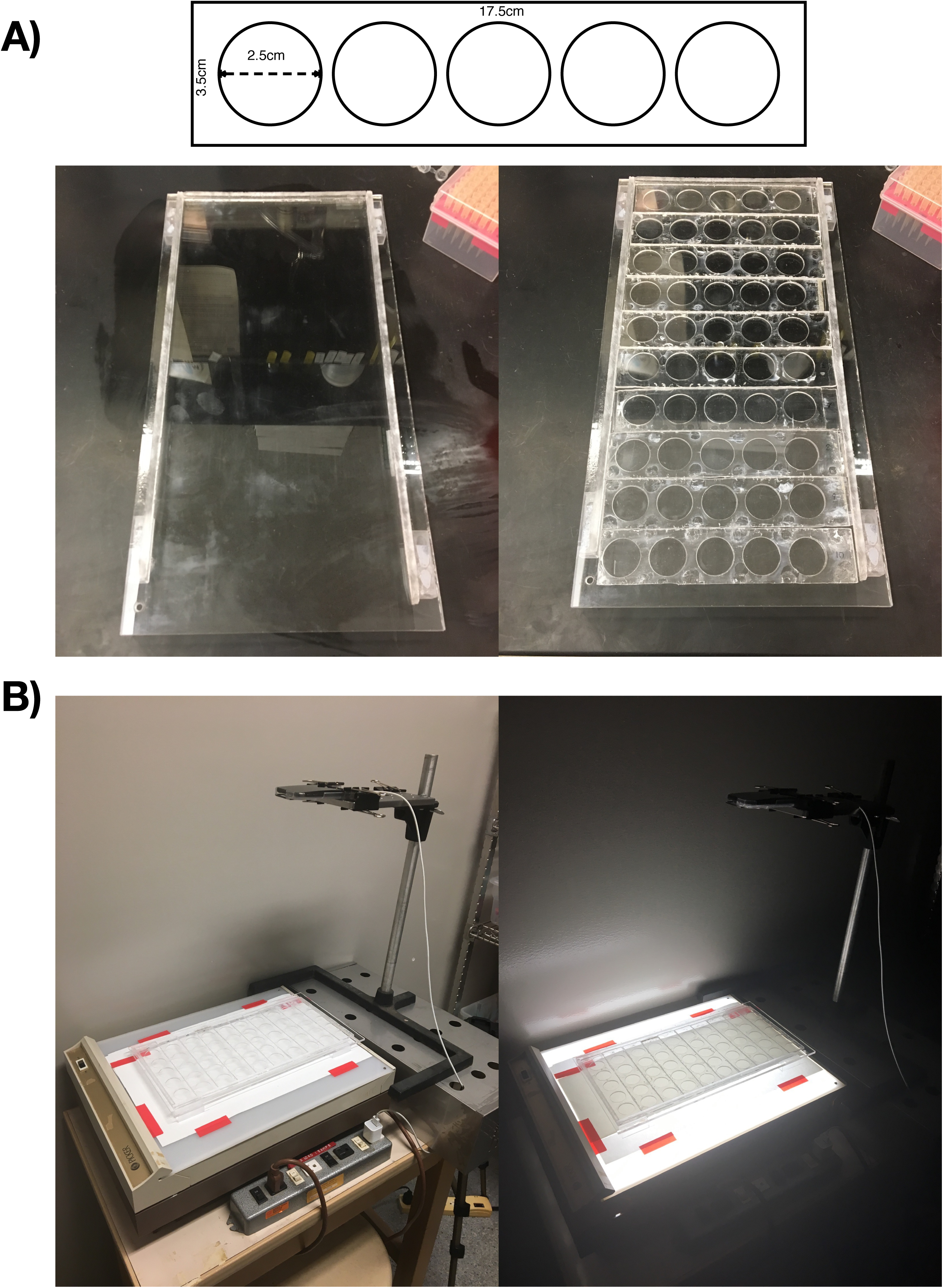
(A) Dimensions of a 5 chamber strip used in the arena. Photo of the base (without strips: left, with strips: right). The strips allow you to customize for the number of chambers needed (5-50). (B) Experimental set up. The base with the loaded chambers is positioned atop A piece of paper taped to the light box to diffuse the light. Recordings are done with the lights in the room off and the light box on to maximize contrast with the fly.

### Fly Husbandry

Crosses were set up in bottles to obtain desired genotypes. Four different wild-type genotypes were obtained by crossing male w^1118^ with UAS-Httex1-Q72-eGFP and UAS-Httex1-Q25-eGFP[21] and female w^1118^ with DJ694 [22] and MHC-Geneswitch [23]. After allowing the parents to mate for 2 days, they were then transferred to a second set of bottles to generate a replicate. The second set of bottles was emptied after two days. Staged flies (0-2 days) were collected under nitrogen anesthesia. A minimum of 4 sample vials for each sex of each genotype containing a minimum of 25 flies were collected. This was repeated for the replicate crosses. Flies were allowed to recover for at least 1 day before being recorded. Flies were maintained on standard fly food (0.01% molasses, 8.2% cornmeal, 3.4% yeast, 0.94% agar, 0.18% benzoic acid, 0.66% propionic acid) at 23-26°C for the duration of the experiment.

### Experimental Setup

A chamber strip was positioned such that the first chamber was circle side down over top of the loading hole of the base. Flies were aspirated from the vial. Once one fly was loaded into the chamber through the loading hole the arena strip is slid such that the second chamber was over top of the loading hole, effectively preventing the fly from escaping. Once all 5 flies in a strip were loaded the strip was slid into place using the guides on the base.

The process was repeated until all the chambers were loaded. The base is then placed atop of a light box, the legs provide an air insulation layer that prevents heating from the light bulbs. A piece of white paper is placed on the light box to diffuse the light. An iPod touch was clamped in a fixed position 40 cm above the light box to capture a single video file encompassing all 50 chambers. Video capturing was performed at 23-26°C with the light table on and the lights in the room off (Figure 1B).

Flies were acclimatized for 1 minute before beginning recording (3 min at 30 frames per second). After recording, flies were aspirated from the arenas out through the hole and returned to their vials. In a given video each genotype was measured in triplicate. Flies were aspirated from one of the four sample vials that were collected, alternating the vial at each recorded time point. The positions of the genotypes in the arena were rearranged each day a recording was done to eliminate bias coming from any specific chamber. Since our genotypes were measured in triplicate, the genotypes shifted by 3 for each time point. Therefore, the number of possible arena configurations is equal to the number of genotypes tested. Supplemental Table 3 shows an example of the various arena configurations for a 16-genotype experiment.

### Software and Files Requirement

Download FlyTracker [20] from: http://www.vision.caltech.edu/Tools/FlyTracker/download.html. Download the DataExtractionScripts (Supplemental Information) and copy and paste each script (Data Extraction Script and Data Compilation Script) in MatLab. The Data Extraction Script is annotated (in green) and includes instructions (in orange) to replace parameters (in red) to accommodate any experimental setup. The Data Extraction Script generates spreadsheet files (.xls and .csv) which, once grouped in a single folder can be processed by the Data Compilation Script to obtain an .xls file combining all experimental data for further analysis, statistical processing and graphing. A custom Arena Configuration table will need to be created (similar to Supplemental Table 3) with the configurations in columns and the chamber numbers in rows. This table can be created in a spreadsheet software and copied into MatLab.

### FlyTracker Tracking

FlyTracker requires the video files to be stored in a folder. We organized our video files in folders by date. The path was set in MatLab for FlyTracker. The ‘tracker.m’ script was run. Specifications for video length, frame rate, and processing options can be selected in the FlyTracker interface. In the Calibrator interface the resolution, number of arenas, number of flies per arena, size and position of arenas, and contrast thresholds for detecting the fly were all selected. Once tracked, all tracking results were verified in the Visualizer interface to ensure accurate tracking. For each video file analyzed, FlyTracker outputs ‘feat.mat’ and ‘track.mat’ structure files.

### Data Analysis

The ‘feat.mat’ and ‘track.mat’ files are dragged and dropped into the Workspace in Matlab and the Data Extraction Script is run to generate spreadsheet files (.xls and .csv). The script uses the Arena Configurations table to assign a genotype according to the chamber number and compiles the distance, the number of frames detecting the presence of the fly, velocity and % time spent immobile (calculated from the number of frames where the fly is immobile). The .xls files are then moved to a single folder and the Data Compilation Script is used to compile them into a single .xls file. Using Microsoft Excel, the values obtained for all 3 individuals for each of the parameters (distance, velocity, % time immobile) were averaged to give a single time point respectively for each parameter. This was executed for both replicates.

## Results and Discussion

Our locomotion measuring device is designed to be adaptable to experiments measuring between 1 and 50 flies (Figure 1). The device was tested with 15 flies (3 strips) and up to 48 flies (10 strips). The resolution of HD 1080p cameras is not sufficient to be able to use more than 10 strips without losing the ability to accurately track every individual fly.

Four wild type phenotypes with different genetic backgrounds were recorded for 3 minutes at multiple time points across age (3-63 days) at the same time of the day to avoid circadian effects. The resulting video files are tracked and analyzed in MatLab using FlyTracker and a custom script to calculate the percentage of time spent immobile and compile it with the instantaneous velocity and the total distance. The distance allows to distinguish between flies that moved sporadically versus those that moved consistently the whole duration of the recording, which may influence their velocity.

Figure 2 (Supplemental Table 1) shows the results of two replicates of four male genotypes UAS-Httex1-Q25-eGFP/+, UAS-Httex1-Q72-eGFP/Y, DJ694/+ and MHC/+. The percentage of time spent immobile increases with all genotypes across age in both replicates (blue and orange). On average, the percentage of time spent immobile remains below 20% for all the genotypes up until day 29 and by late life (50-60 days) the percent immobility reaches about 80%. Additionally, the distance traveled and velocity declined with all male genotypes across age in both replicates. On average, the distances traveled begins to decline at day 16 and remains below 10000mm into late life. With the velocity, all genotypes started with an average between 10-15mm/s then at day 24-29 decline to below 10mm/s.

**Fig. 2.**
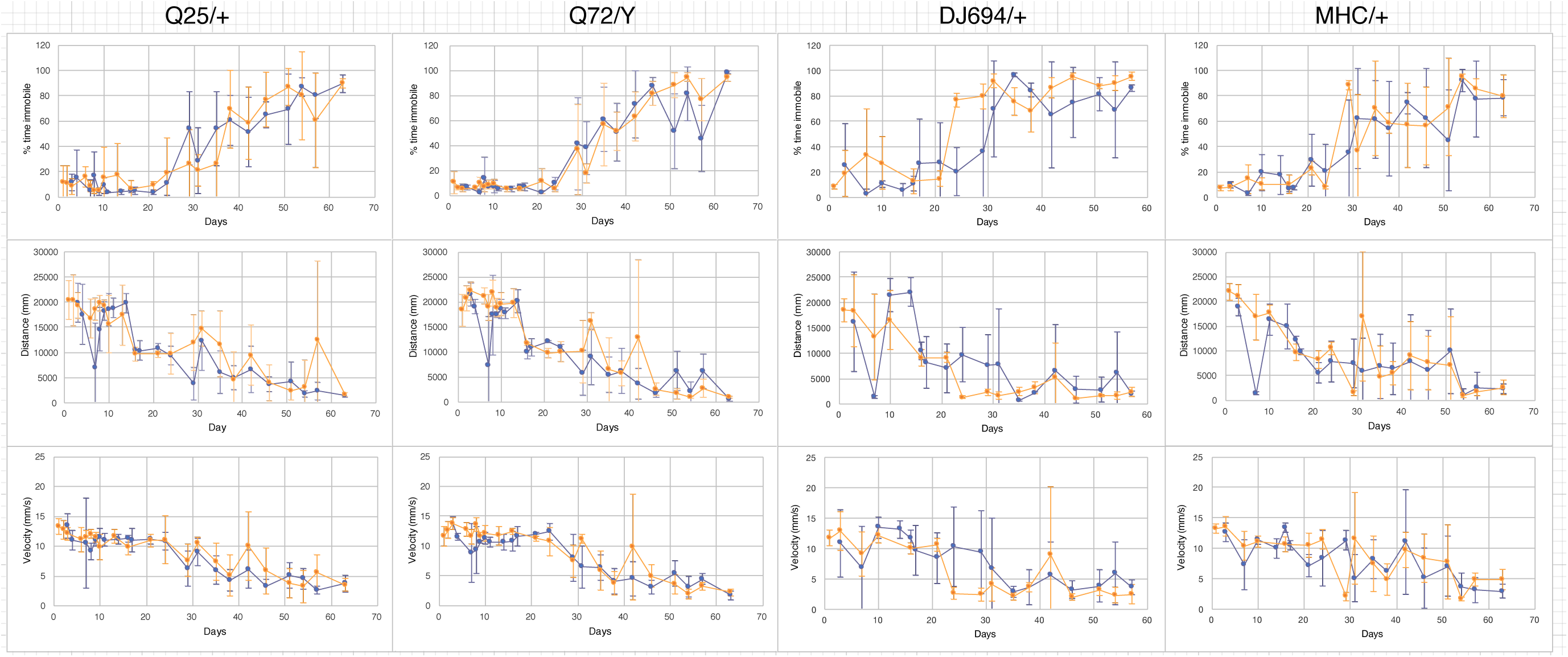
Locomotion of male UAS-Httex1-Q25-eGFP/+,UAS-Httex1-Q72-eGFP/Y, DJ694/+ and MHC/+ flies across age (x axis) as measured by percentage of time spent immobile (top), distance traveled (middle) and velocity (bottom). The blue and orange lines denote each of the independent replicates (each replicate n=3). Error bars represent ±SD.

Figure 3 (Supplemental Table 2) shows the results of two replicates of four female genotypes UAS-Httex1-Q25-eGFP/+, UAS-Httex1-Q72-eGFP/X, DJ694/+ and MHC/+. The percentage of time spent immobile increases with all genotypes across age in both replicates (blue and orange). The female genotypes showed very similar results to the males, on average the percentage of time spent immobile remained below 20% and by day 50-60 it reaches about 80%. Also, the distance traveled and velocity showed similar results to the males, at day 16 the distance declined below 10000mm and the velocity decreased below 10mm/s at day 24-29.

**Fig. 3.**
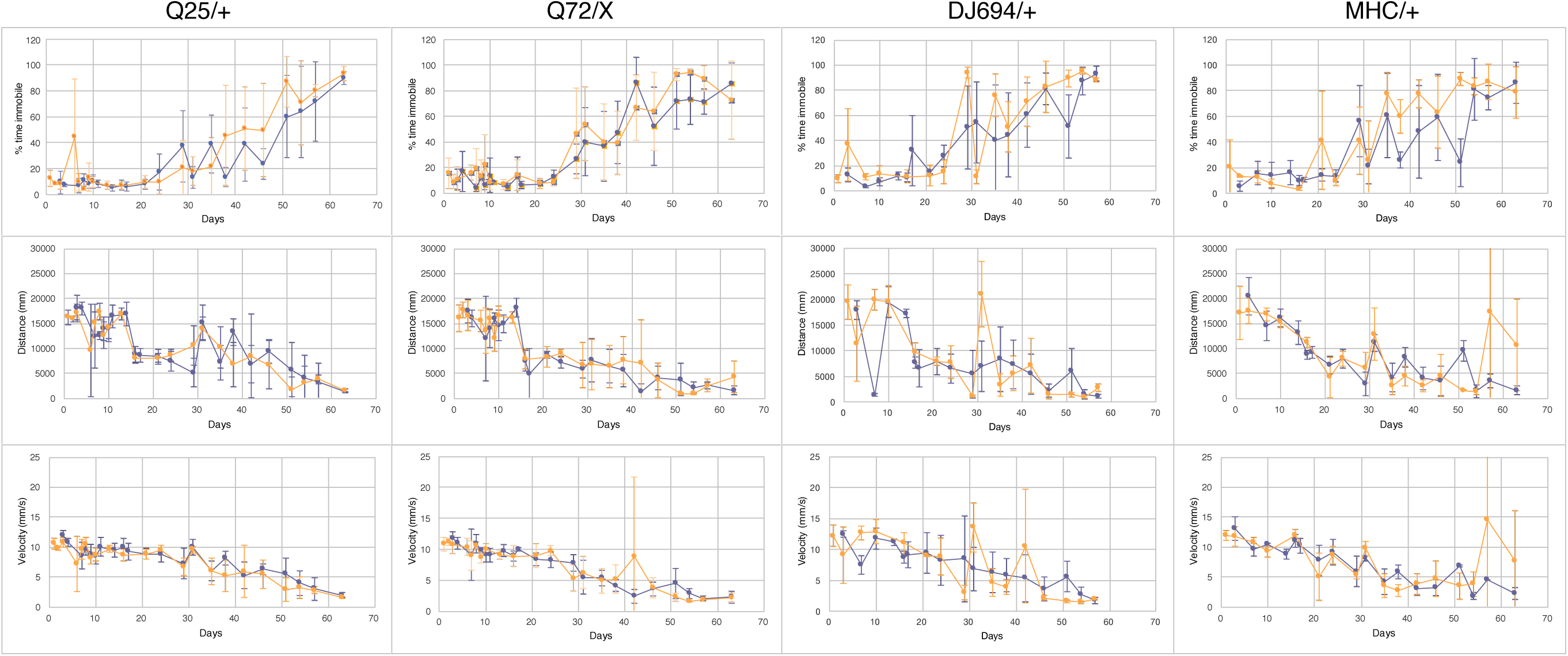
Locomotion of female UAS-Httex1-Q25-eGFP/+, UAS-Httex1-Q72-eGFP/X, DJ694/+ and MHC/+ flies across age (x axis) as measured by percentage of time spent immobile (top), distance traveled (middle) and velocity (bottom). The blue and orange lines denote each of the independent replicates (each replicate n=3). Error bars represent ±SD.

It has been known for a long time that advancing age is correlated with behavioral declines. Declines in negative geotaxis, flight and locomotion have previously been reported with a variety of different experimental approaches [24-30]. The method used in this study yields reproducible results that indeed confirm in multiple genotypes and both sexes that normal aging is associated with a progressive decline in locomotion as of result of increased immobility and reduced velocity.

---

**Supplemental Table 1:**
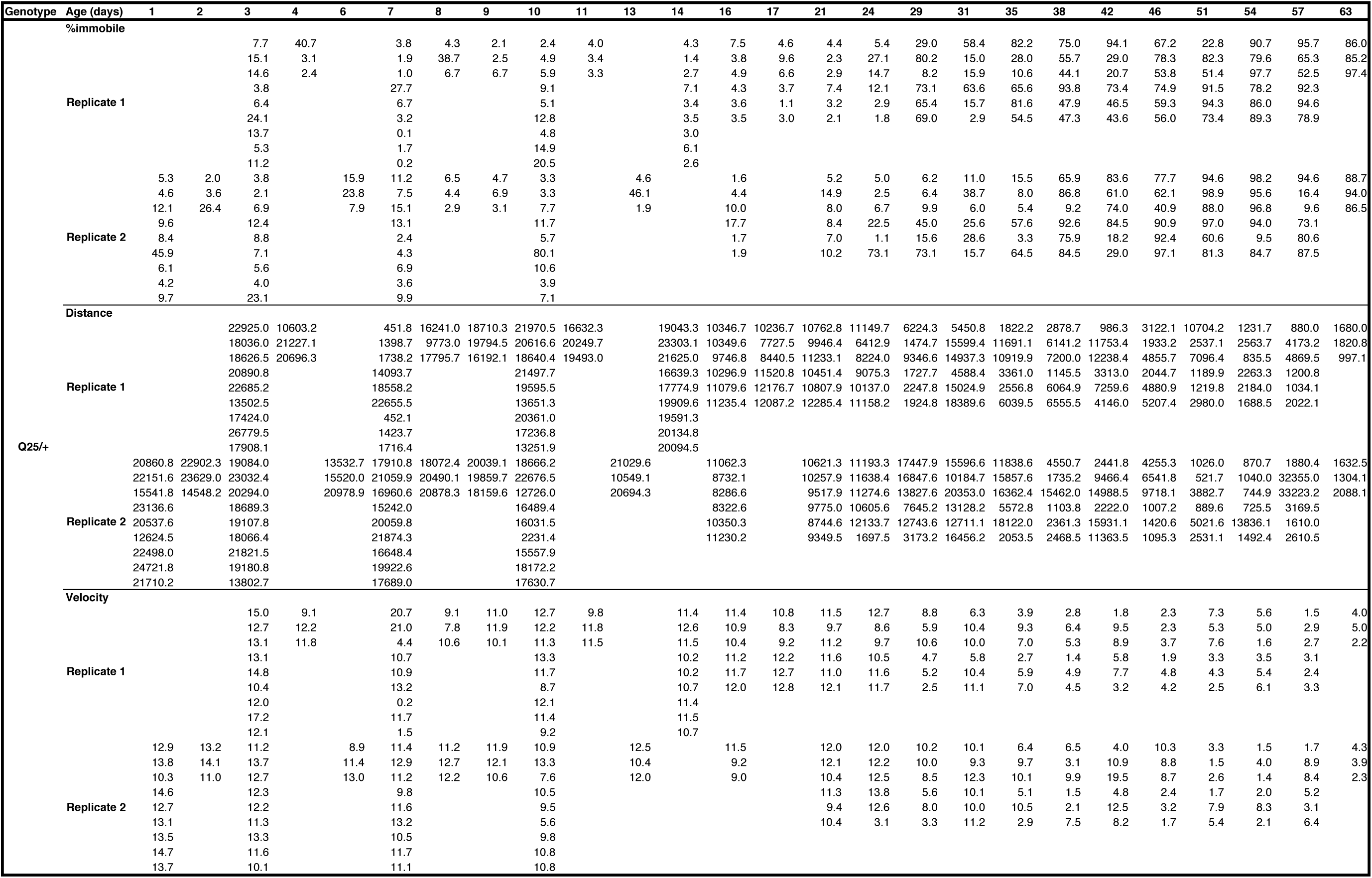
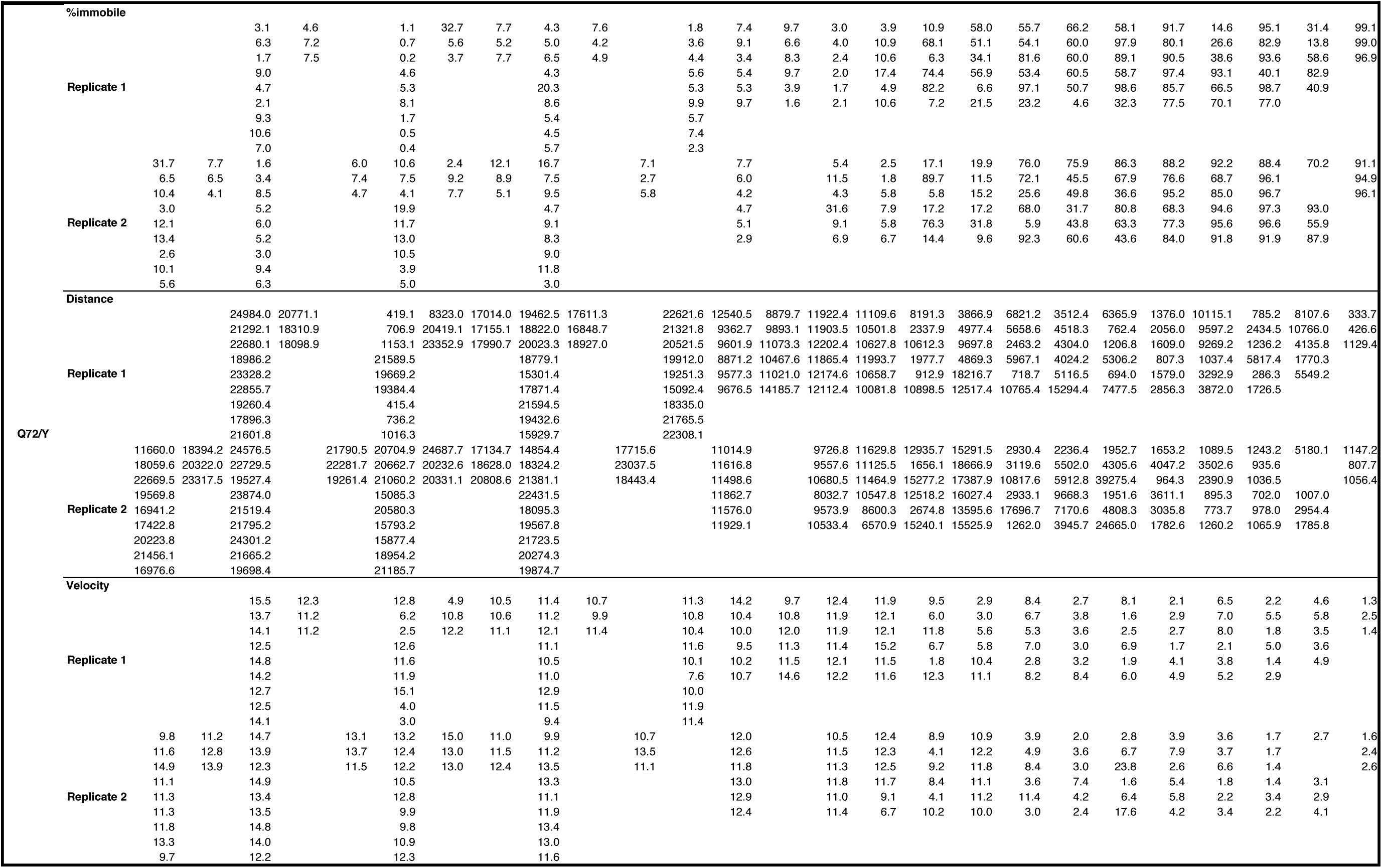
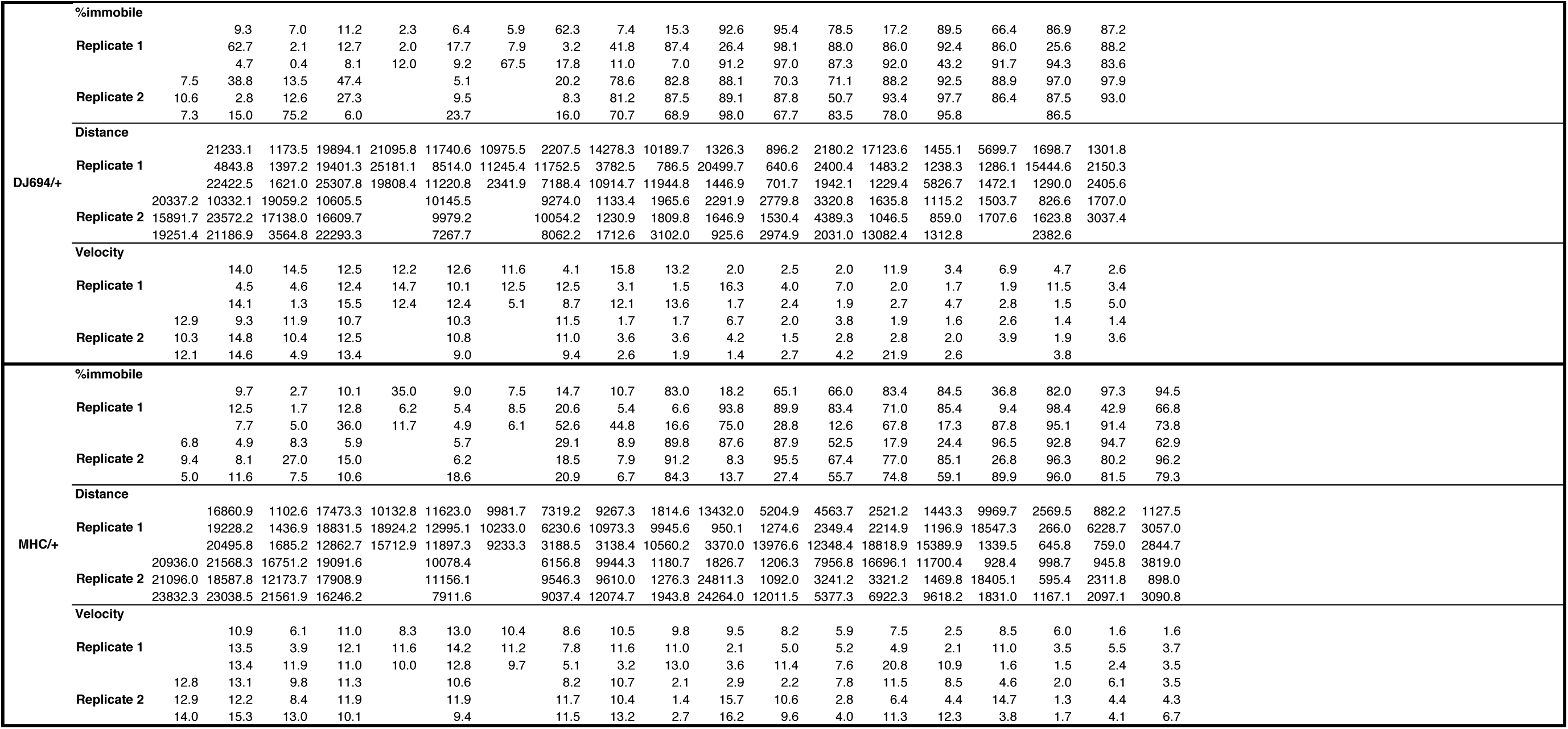
Raw locomotor behaviour data from male genotypes. Each value represents one individual fly that was measured. All individuals of a genotype were averaged for each replicate in the final figures.

**Supplemental Table 2:**
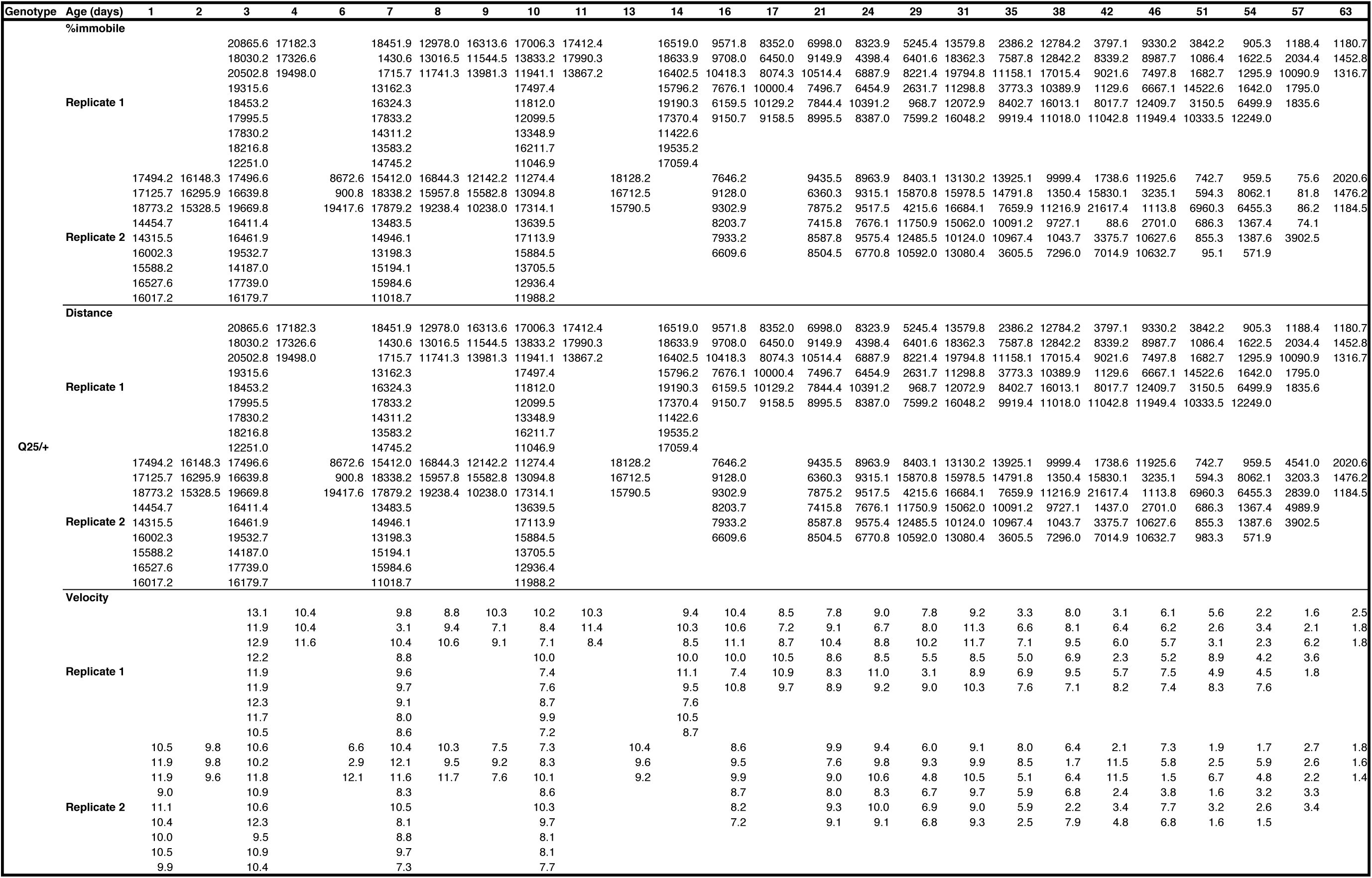
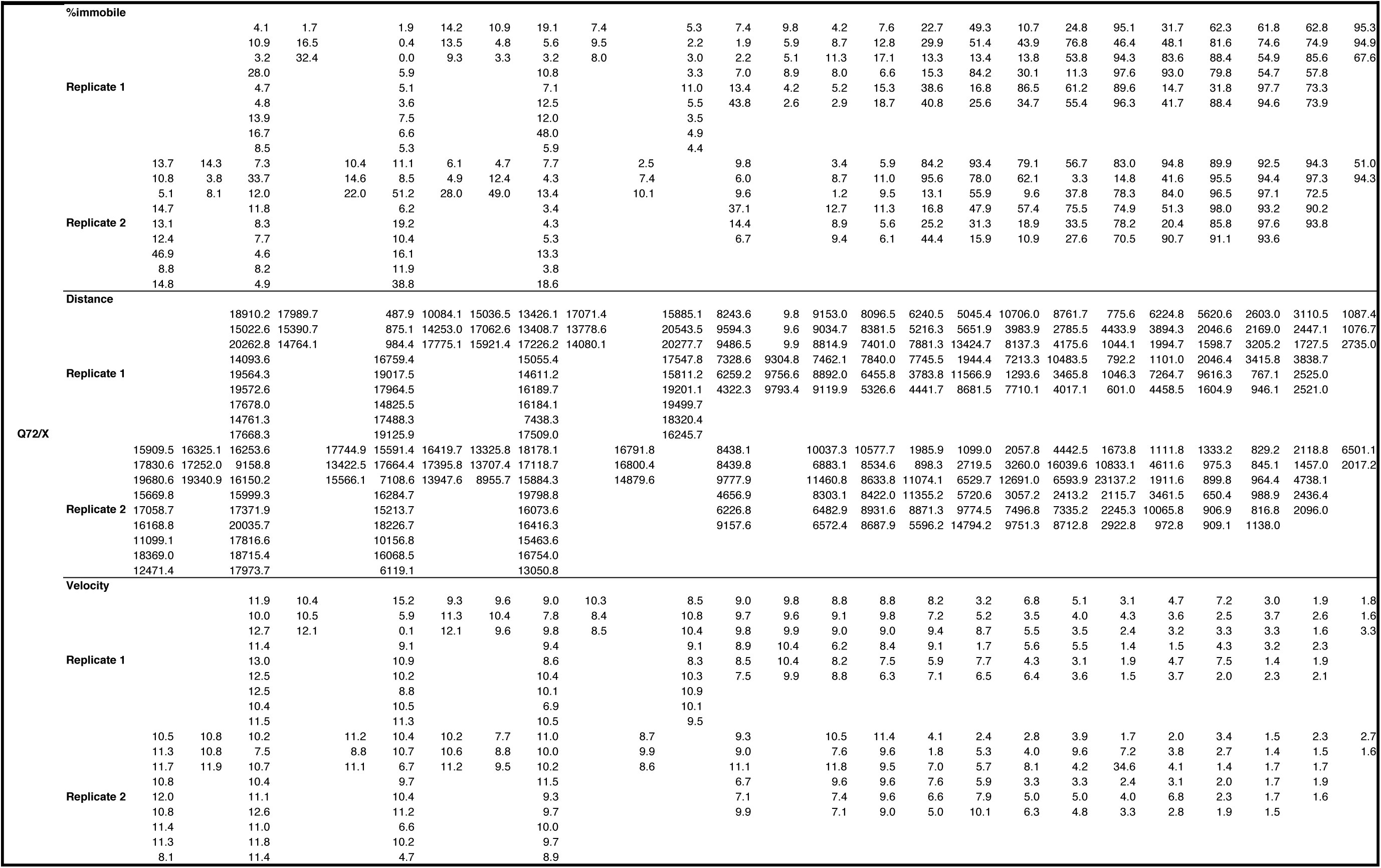
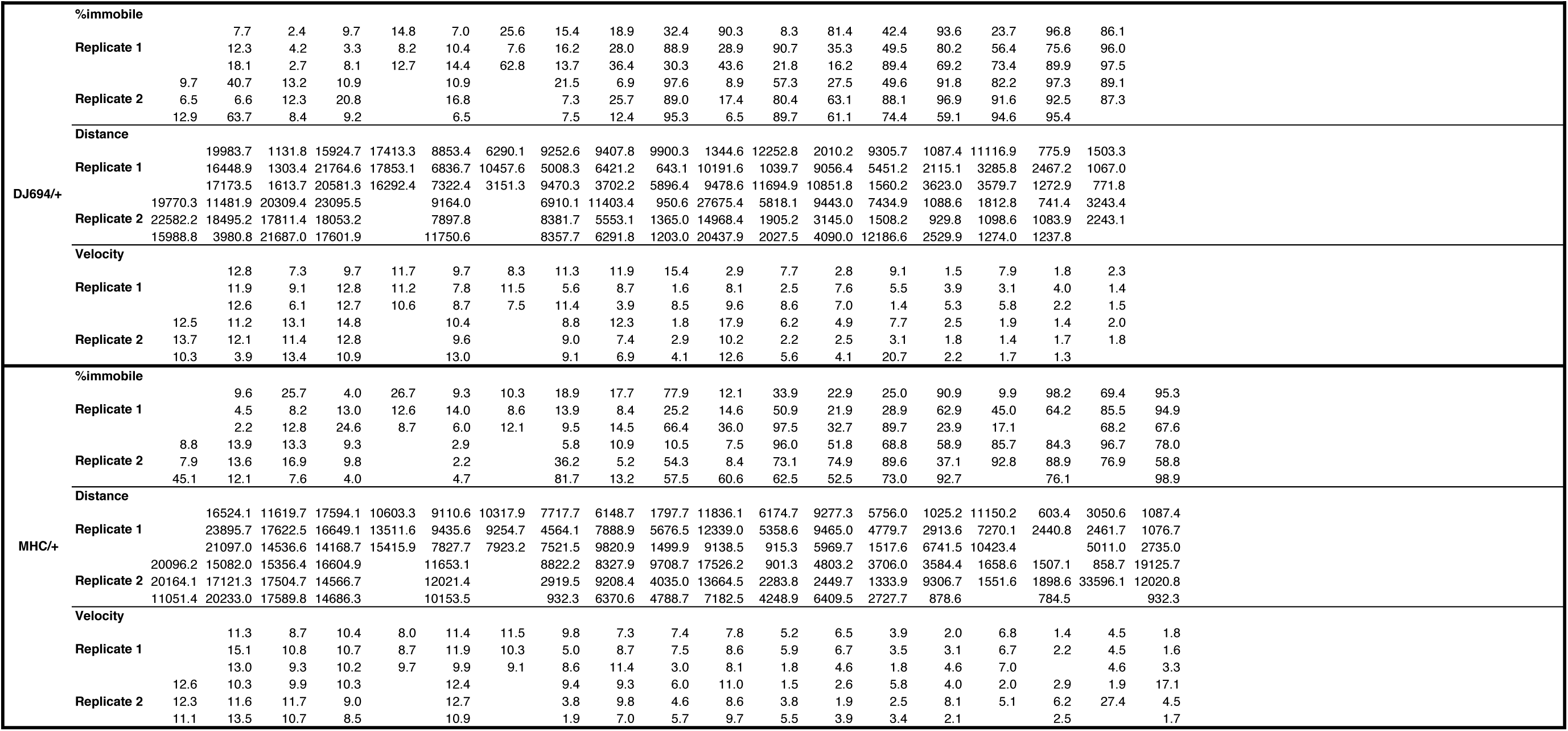
Raw locomotor behaviour data from female genotypes. Each value represents one individual fly that was measured. All individuals of a genotype were averaged for each replicate in the final figures.

**Supplemental Table 3:**
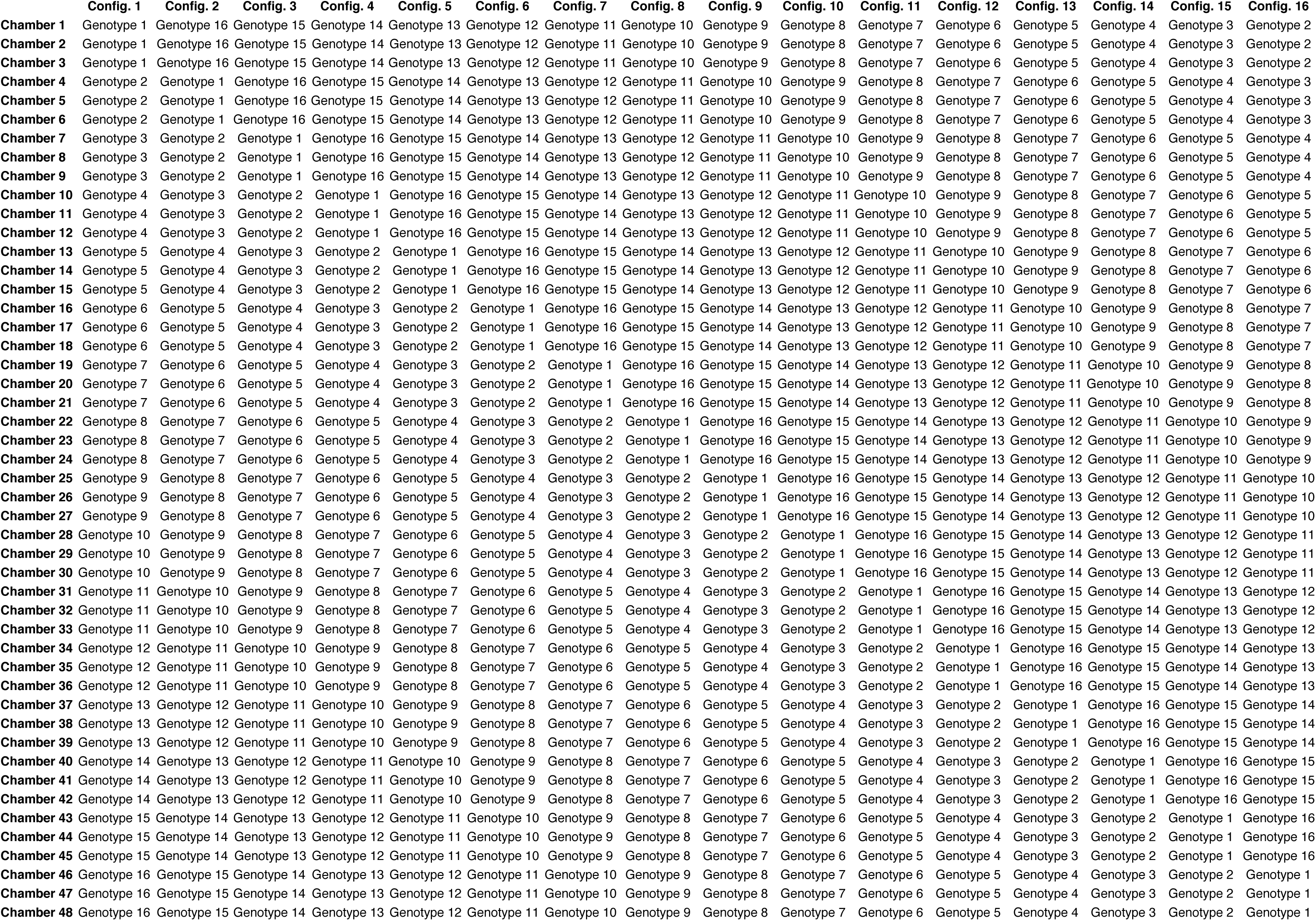
Sample arena configuration table used by the extraction script to identify which genotype is in each chamber based on the configuration that is input. The number of configurations is equal to the number of genotypes tested. The number of chambers required is equal to the number of genotypes tested multiplied by the sample size of each genotype.

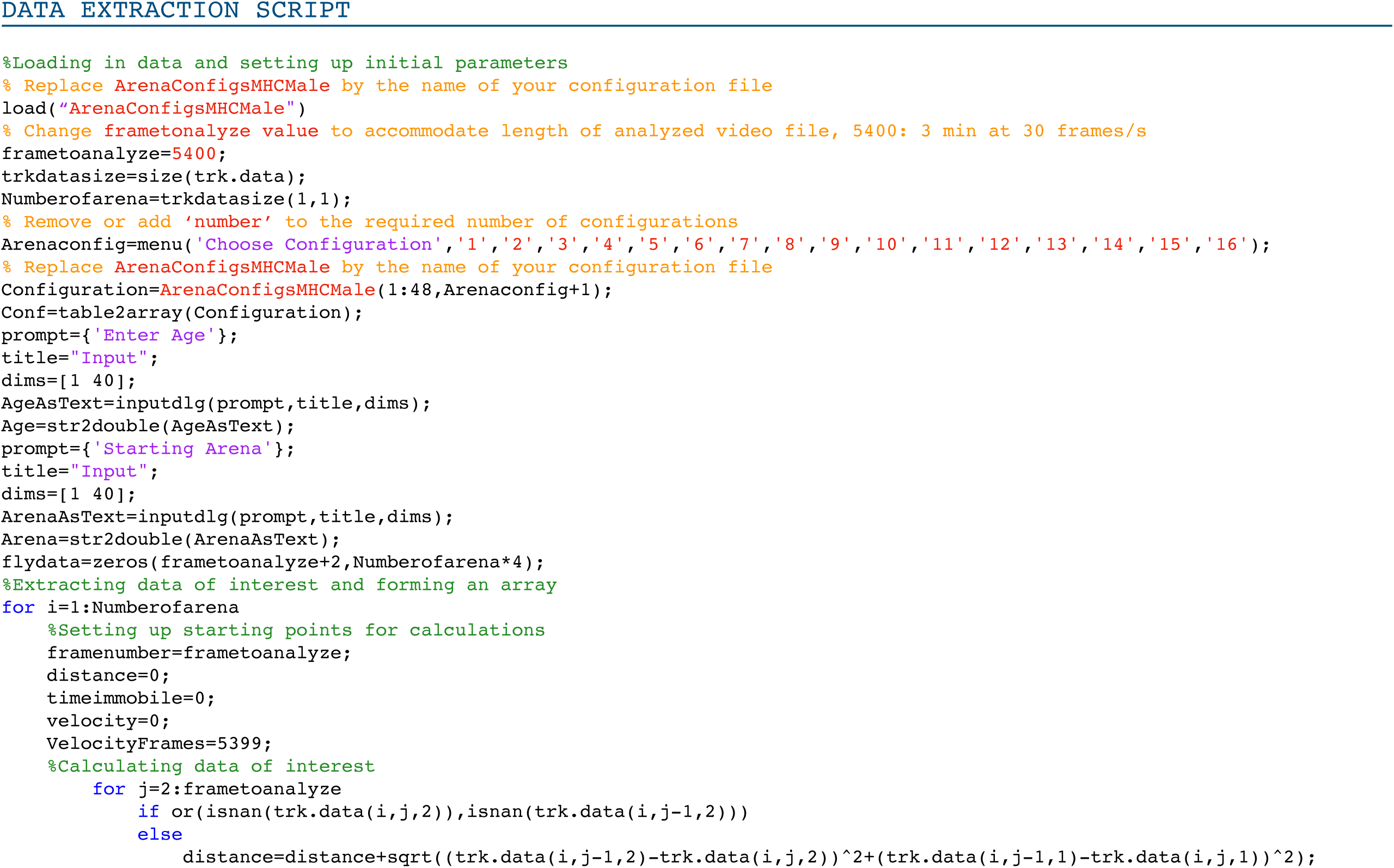
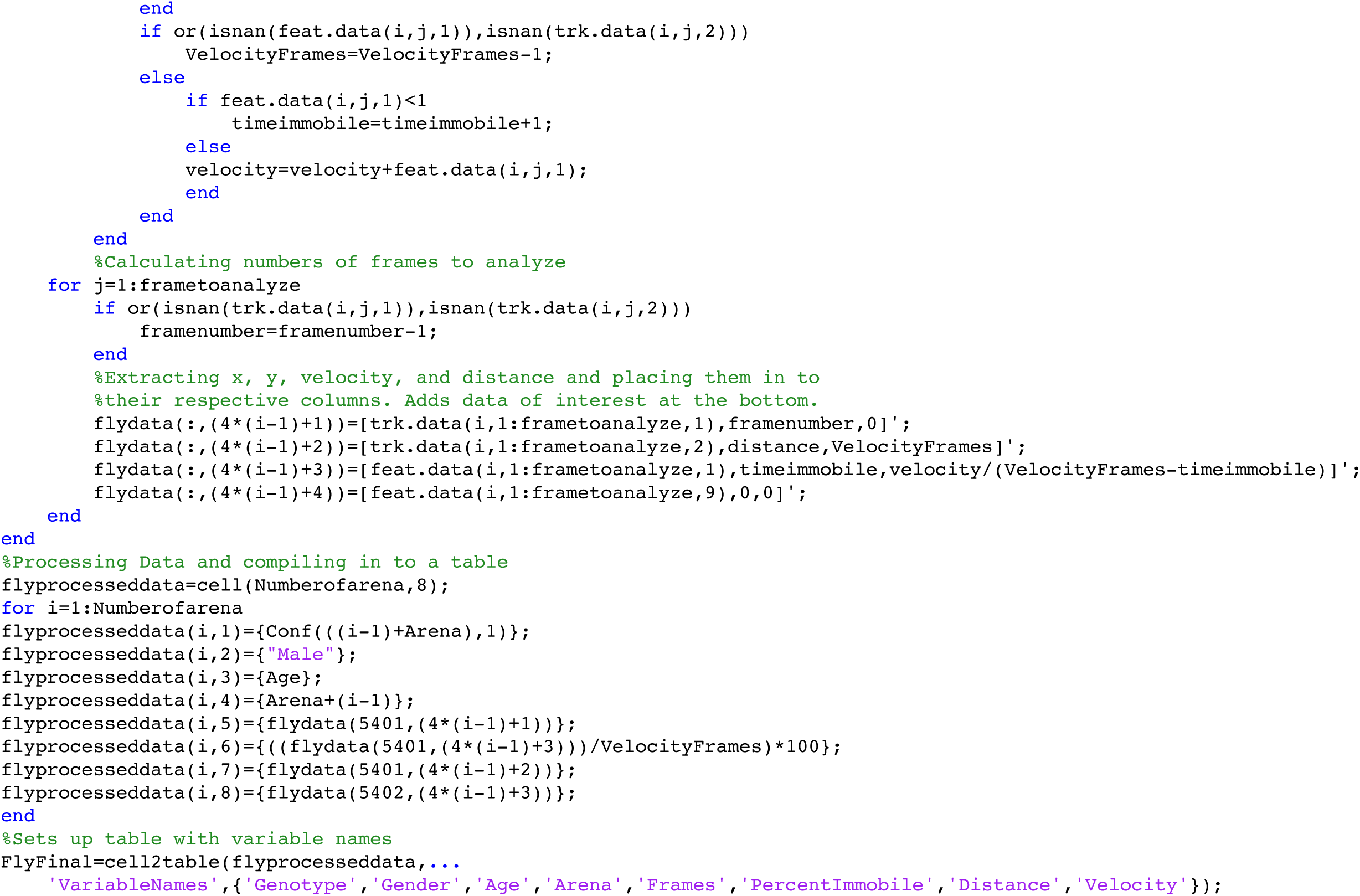
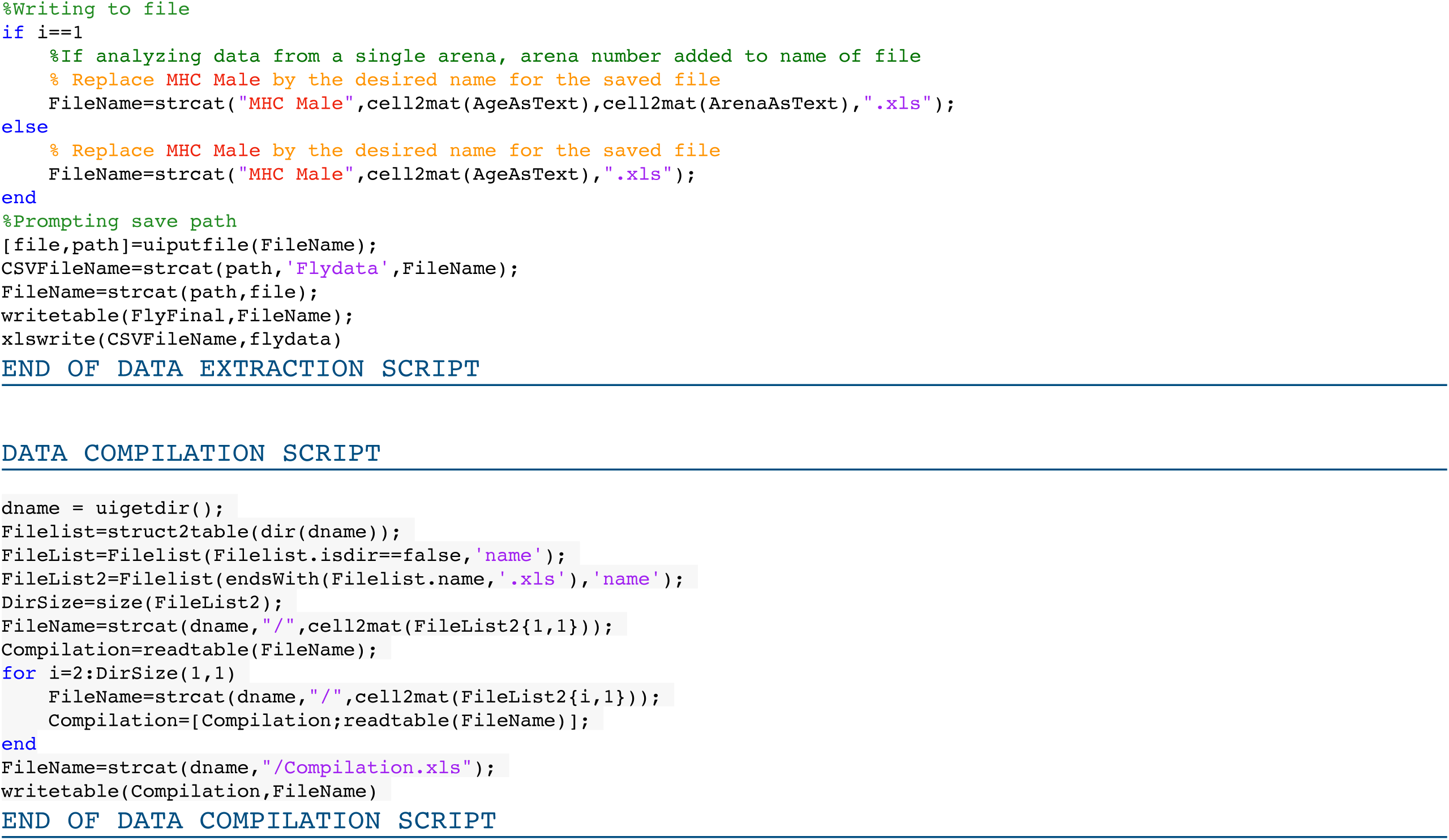

